# Wetland plant evolutionary history influences soil and endophyte microbial community composition

**DOI:** 10.1101/2020.06.22.165738

**Authors:** Marisa B. Szubryt, Kelly Skinner, Edward J. O’Loughlin, Jason Koval, Stephanie M. Greenwald, Sarah M. Owens, Kenneth M. Kemner, Pamela B. Weisenhorn

## Abstract

Methane is a microbially derived greenhouse gas whose emissions are highly variable throughout wetland ecosystems. Differences in plant community composition account for some of this variability, suggesting an influence of plant species on microbial community structure and function in these ecosystems. Given that closely related plant species have similar morphological and biochemical features, we hypothesize that plant evolutionary history is related to differences in microbial community composition. To examine species-specific patterns in microbiomes, we selected five monoculture-forming wetland plant species based on the evolutionary distances among them. We detected significant differences in microbial communities between sample types (unvegetated soil, bulk soil, rhizosphere soil, internal root tissues, and internal leaf tissues) associated with these plant species based on 16S relative abundances. We additionally found that differences in plant evolutionary history were correlated with variation in microbial communities across plant species within each sample type. Using qPCR, we observed substantial differences in overall methanogen and methanotroph population sizes between plant species and sample types. Methanogens tended to be most abundant in rhizosphere soils while methanotrophs were the most abundant in roots. Given that microbes influence methane flux and that plants affect methanogen and methanotroph populations, plant species contribute to variable degrees of methane emissions. Incorporating the influence of plant evolutionary history into future modeling efforts may improve predictions of wetland methane emission since microbial community differences correlate with differences in plant evolutionary history.

## Introduction

Wetlands are globally important ecosystems which are responsible for 20-39% of global methane emissions (IPCC 2007). Methane is largely a microbially derived greenhouse gas that is 25 times more potent than carbon dioxide when considered over a 100-year horizon (Boucher *et al*. 2009) and is an important component of global climate change (Cao *et al*. 1998). Wetlands contain heterogeneous soils with spatial and temporal variation in oxidation-reduction (redox) potentials that affect microbial communities as well as controlling microbial metabolic processes (Conrad 1996; Megonigal 2004; Pett-Ridge and Firestone 2005), including the production and consumption of methane. The presence of abundant anoxic soil habitats within wetlands allow for the growth of methane-producing microbes known as methanogens.

Methane emissions from wetlands are a highly variable and difficult to accurately interpolate component of global climate change models (Bartlett *et al*. 1989; Ringeval *et al*. 2010; Yavitt and Knapp 1998). Temporal and spatial heterogeneity in methane emissions can be partially explained by differences in plant activity (Wang and Han 2005) and species composition (Chanton *et al*. 1989; Grünfeld and Brix 1999). Thus, the influence of wetland plants on methane emissions may be a consequence of their effect on methanogens and methane-consuming microbes known as methanotrophs. Wetland soil microbial communities may be affected by plant species-specific variation in the abundance and form of organic matter available or from differences in radial oxygen loss (Fechner-Levy and Hemond 1996; Grünfeld and Brix 1999; Hackstein and Steinbüchel 2010).

Many wetland plants have evolved a spongy tissue known as aerenchyma (Evans 2004). Aerenchyma permits aerobic root respiration under anoxic soil conditions by facilitating oxygen transport from leaf to root tissues (Visser *et al*. 2000). Aerenchyma can also function as a conduit for methane release from soil to the atmosphere (Colmer 2003; Yavitt and Knapp 1998), as gases can diffuse 10,000 times faster through gas-filled plant tissues than through water filled soil pores. Conductive flow rates through aerenchyma can vary drastically by plant species (Brix *et al*. 1992). For example, *Phragmites australis* and *Typha domingensis* have rates two orders of magnitude greater than other wetland plants (Brix *et al*. 1992; Konnerup *et al*. 2011). Differences in flow rates may result from variation in the total volume or type of aerenchyma (Jung *et al*. 2008). Plants species which are close evolutionary relatives and occupy similar habitats frequently have similar aerenchyma morphology (Jung *et al*. 2008; Visser *et al*. 2000), thus may similarly influence oxygen diffusion through roots and into adjacent soil (Visser *et al*. 2000). This radial oxygen loss can subsequently change soil redox potentials (Colmer 2003), influencing microbial communities (Conrad 1996; Nikolausz *et al*. 2008) by altering the thermodynamic favorability of specific microbial metabolic reactions.

Wetland plant species may also influence microbial community structure and function through the production of organic matter via senesced tissues (Findlay *et al*. 2002; Newell *et al*. 1995) and root exudates (Bertin *et al*. 2003). Differences in plant productivity and biomass (Clarke and Baldwin 2002) have been shown to correlate with differences in leaf litter chemistry (Hobbie 1992) and root exudate production (Valé *et al*. 2005) in other ecosystems. Further, these differences in organic matter inputs may be related to plant evolutionary history (Reich *et al*. 2003). Wetland plants may produce lineage-specific root exudates to recruit particular microbial symbionts as occurs in other ecosystems (Bais *et al*. 2006; Haichar *et al*. 2014). Such interspecific differences in the quality and quantity of aboveground plant litter (Davis and van der Valk 1978) and root exudates (Bais *et al*. 2006), particularly low-molecular weight compounds, are known to affect adjacent soil microbial communities by altering resource availability (Conrad 1996; Westermann *et al*.1989).

In addition to their effects on soil-dwelling microbes, plants are hosts to their own endophytic microbial communities. Unique endophyte communities may reside in specific plant tissues (Bai *et al*. 2015; Hardoim *et al*. 2015; Llirós *et al*. 2014; Ma *et al*. 2013) or specific host taxa (Li *et al*. 2013; Winston *et al*. 2014). Endophytic microbial community composition is influenced both by environmental factors (Hardoim *et al*. 2015; Ma *et al*. 2013) and conditions within host tissues (Espinosa-Garcia and Langenheim 1990; Sun *et al*. 2008; Wemheuer *et al*. 2017). Plant tissues may provide more stable conditions relative to stochastic soil environments (Hallmann *et al*. 1999; Rosenblueth and Martínez-Romero 2006). However, endophytic microbial communities can also be directly influenced by the host immune system (Bulgarelli *et al*. 2012), biochemistry (Hardoim *et al*. 2008; Mengoni *et al*. 2010), and additional stressors (Helander *et al*. 1993). Evolutionarily conserved physiological and biochemical differences among plant species (Bohlmann *et al*. 1998; Giannasi 1978; Wink 2003) likely contribute to differences in endophytic microbial communities.

This study examines whether wetland plant evolutionary history influences the composition of soil and endophyte microbial communities with an emphasis on methanogen and methanotroph populations. Specifically, we hypothesized that more closely related plant species would have more similar microbial communities in each of four sample types along an anoxic to oxic gradient: bulk soil, rhizosphere soil, root tissue, and leaf tissue. We expected that oxygen-sensitive methanogen population sizes would decrease along this gradient from bulk soil to leaf tissue and that oxygen-dependent methanotroph population sizes would follow the inverse pattern, increasing from bulk soil to leaf tissue. This study illuminates the relationship between host evolutionary distances and microbial community dissimilarities within sample types. Further, we highlight variability in methanogen and methanotroph populations among plant species and sample type.

## Materials and Methods

### Plant selection and sample collection

Five monoculture-forming wetland plant species in the grass order Poales were selected to examine the influence of individual species on soil and endophyte microbial communities. Species sampled include representatives of the cattail (*Sparganium eurycarpum* Engelm., *Typha* × *glauca* Godr.), true grass (*Phalaris arundinacea* L., *Phragmites australis* (Cav.) Trin. ex Steud.), and sedge (*Bolboschoenus fluviatilis* (Torr.) Soják) families. The cattail family species have been reported as obligate wetland plants with efficient aerenchyma and high flooding tolerance that are typically poor competitors under drier conditions (Day *et al*. 1988; Sulman *et al*. 2013; Tuchman *et al*. 2009; USDA 2017). The true grass and sedge species have been defined as facultative wetland plants with aerenchyma and occupy less intensely and less frequently flooded habitats (Brix *et al*. 1992; Davis and van der Valk 1978; Kercher and Zedler 2004; Wetzel and van der Valk 1998). The evolutionary history and phylogenetic distances of these taxa have been well studied (Givnish *et al*. 2010).

Three replicate individuals of each species were sampled at one of two sites: wetland one, where *B. fluviatilis, P. australis*, and *T*. × *glauca* were located, and wetland two, where *P. arundinacea and S. eurycarpum* were located. These wetlands were within 500 m of each other and were hydrologically connected. We collected four sample types (bulk soil, rhizosphere soil, root tissue, leaf tissue) from each individual to test the effects of plant species on soil and endophytic microbial communities. Additionally, we collected three replicates of unvegetated soil at least one meter away from visible plant shoots at each site. Unvegetated and bulk soils were collected using a 16 cm deep, 2 cm diameter PVC core. Bulk soil was collected between 10 cm and 30 cm away from plant shoots. Rhizosphere soil was collected by digging up an intact plant root system and collecting soil adhered to the roots. Soil samples were placed on dry ice in the field. Each root sample consisted of four randomly selected root sections collected from a single plant. A randomly selected, mature, non-senescent leaf from each plant was also collected.

### Sample Preparation and Sequencing

Roots and leaves were initially rinsed with tap water followed by deionized water. We then surface sterilized 1 cm-portions by placing them in a 95% ethanol solution for 30 s. Tissues were then transferred to a 10% bleach solution for 3 min followed by a 70% ethanol solution for 3 min to remove DNA from the outer surface (Cao *et al*. 2004). Soil cores from unvegetated and bulk soil were homogenized and sampled to avoid contamination with root material. Rhizosphere soil was gently scraped from the surface of roots directly into bead-beating tubes.

We used the DNeasy PowerSoil Kit (Qiagen) to extract microbial gDNA from 250 mg wet weight of each sample. The protocol for this kit was followed, with the addition of an initial ten-minute heating step at 65°C in accordance with the Earth Microbiome Project DNA extraction protocol (Gilbert *et al*. 2014; Marotz *et al*. 2017). Extracted DNA was stored in a −80°C freezer until sequencing. The 515F (GTGYCAGCMGCCGCGGTAA) and 806R (GGACTACNVGGGTWTCTAAT) primers were used to amplify the V4 region of the prokaryotic 16S rRNA gene (Bergmann *et al*. 2011; Caporaso *et al*. 2012; Lundberg *et al*. 2013). Peptide nucleic acid (PNA)-clamps were included in the master mix for root and leaf tissues to minimize plant mitochondrion (GGCAAGTGTTCTTCGG) and chloroplast (GGCTCAACCCTGGACAG) amplification (Lundberg *et al*. 2013). PCR amplification consisted of 45 s at 95°C, followed by 34 cycles of 95°C for 15 s, 78°C for 10 s, 60°C for 30 s, and extension 72°C for 30 s. The program terminated with a cooldown to 4°C. The Environmental Sample Preparation and Sequencing Facility at Argonne National Laboratory sequenced the amplicon libraries using the Illumina Mi-Seq platform with V3 chemistry.

Quantitative PCR (qPCR) was performed in triplicate to determine methanogen and methanotroph abundances by amplifying the *mcrA* (Steinberg and Regan 2009) and *pmoA* genes (Kolb *et al*. 2003), respectively. The primers *mcrA* mlasFW (GGTGGT GTMGGDTTCACMCARTA) and *mcrA*RV (CGTTCATBGCGTAGTTVGGRTAGT) were used to determine the gene copy number and abundance of methanogens in each sample. The primers *pmoA* (MTOT) A189 FW (GGNGACTGGGACTTCTGG) and *pmoA* (MTOT) mb661RV (CCGGMGCAACGTCYTTACC) were used to determine gene copy number of methanotrophs. Unlike mcrA which has a single copy per cell, two copies of the *pmoA* gene are frequently present (Kolb *et al* 2003). DNA was quantified using PicoGreen. Two microliters of DNA extract, containing 0.5 ng/μL to 10.0 ng/μL DNA, were added to each well containing master mix. Samples were amplified on a Roche LightCycler 480 instrument. The *pmoA* gene was amplified at 95°C for 5 min followed by 50 cycles of 95°C for 20 s, 65°C for 20 s, and 72°C for 45 s. The *mcrA* gene was amplified at 95°C for 5 min followed by 50 cycles of 95°C for 30 s, 55°C for 45 s, and 72°C for 30 s. The *Methanobacterium thermautotrophicum mcrA* gBlock was used as a quantification standard. The *Methylococcus capsulatus pmoA* gBlock was used for methanotroph quantification. The gBlocks gene copy curve ranged from 10^1^ to 10^8^ copies/μL.

### Plant phylogenetic distance

Internal transcribed spacer data from the five study plants (*B. fluviatilis, P. arundinacea, P. australis, S. eurycarpum*, and *T. latifolia*, the latter a parent species of *T*. × *glauca)* were downloaded from GenBank, along with the outgroup *Sabal minor*. Sequences were aligned in SeaView (Galtier *et al*. 1996) using Muscle (Edgar 2004). Parsimony heuristic phylogenetic analyses were conducted in PAUP v4.0 (Swofford 2002) with *S. minor* as an outgroup. Patristic phylogenetic distances were calculated from total branch lengths between plant species (SFig. 1).

**Figure 1.**
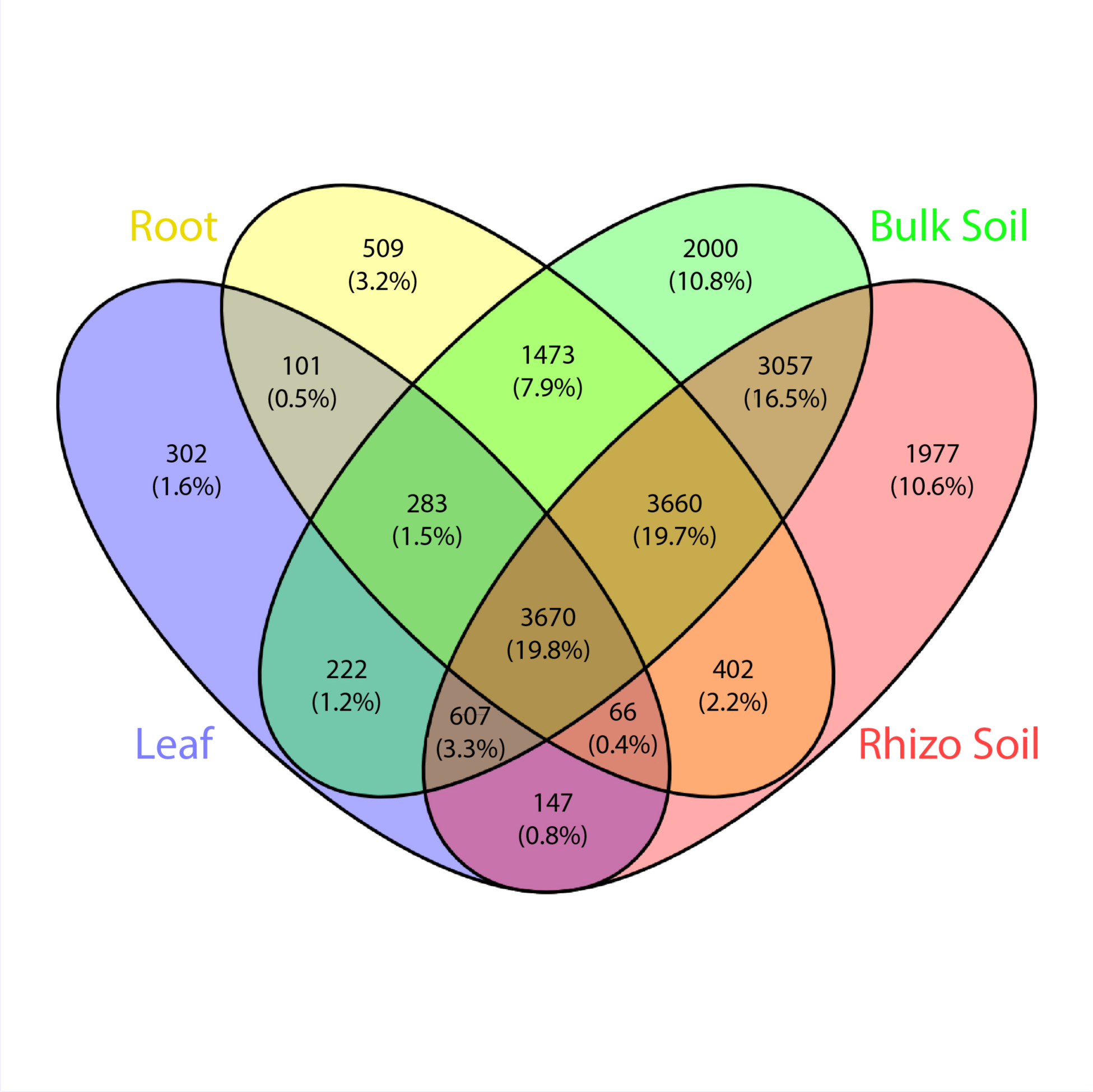
Number of microbial OTUs found in each sample type or shared among sample types. Sample types and combinations are indicated with different colors.

### Data Analysis

Sequence data were processed using QIIME (Caporaso *et al*. 2010) and UPARSE (Edgar 2013). Reads were paired and demultiplexed using QIIME while unpaired reads, reads with barcode errors, reads of incorrect length, and sequences with a quality below q20 were removed. Next, sequences were dereplicated, sorted by size, had singletons removed, and clustered *de novo* using the UPARSE algorithm (Edgar 2013) with operational taxonomic units (OTUs) assigned at 97% sequence identity. All remaining data analyses were conducted using R v3.4.3 (R Core Team 2013).

Shannon diversity values were calculated for each sample using the vegan package (Oksanen *et al*. 2007). All samples contained greater than 10,000 reads and were therefore included in diversity analyses. ANOVA and Tukey’s honest significant difference (HSD) tests were used to determine differences in diversity among sample types, plant species, and sites (de Mendiburu 2017). Relative abundance data were used for all compositional analyses to account for differences in sampling depth. ANOSIM and PERMANOVA analyses were conducted to determine the relative effects of sample type, plant species, and wetland site using the base and vegan packages in R, respectively. These results were visualized using non-metric multidimensional scaling (NMDS) ordinations produced using vegan. Analyses and NMDS visualization were performed for all sample types and each sample type separately to examine the effects of sample type, plant species, and site. To evaluate the influence of plant evolutionary history on the composition of microbial communities, Mantel tests were used to compare the evolutionary distances among plant species with the Bray-Curtis distances among microbial communities and were performed for each sample type individually. Differences in methanogen and methanotroph population sizes among sample types and plant species were examined using ANOVA and Tukey’s honest significant difference test. For root and leaf samples, gene copy number per ng of DNA was reported. Gene copy number for soil samples were reported relative to unvegetated controls by site to account for background variation in methanogen and methanotroph population sizes between the sites.

## Results

### Community Composition

Overall, microbial communities were most strongly influenced by sample type. Sample type explained 52% of the observed variation across all samples, and microbial communities clustered distinctly by sample type (ANOSIM p=0.001, R = 0.86) (SFig. 2). Conversely, plant species explained 18% of the observed variation across all samples (p=0.001) while wetland site accounted for 7% of the observed variation (p=0.001). Alpha diversity was significantly different among sample types (p<0.0001). Rhizosphere communities had higher Shannon’s diversity than bulk soil communities; bulk communities had higher diversity than root communities, which had higher diversity than leaf communities. There were 13,586 total OTUs in bulk soil samples, 14,972 OTUs in rhizosphere soil samples, 10,254 OTUs in root samples, and 5,398 OTUs in leaf samples (Fig. 1). Bulk and rhizosphere soil samples had high numbers of unique OTUs (1,977 and 2,000, respectively) compared to either root (599) or leaf (302) sample types. Nearly 40% of all OTUs were shared across all sample types (3,670 or 19.8% of total), with an additional 3,660 shared among roots and soil types (3,660).

**Figure 2.**
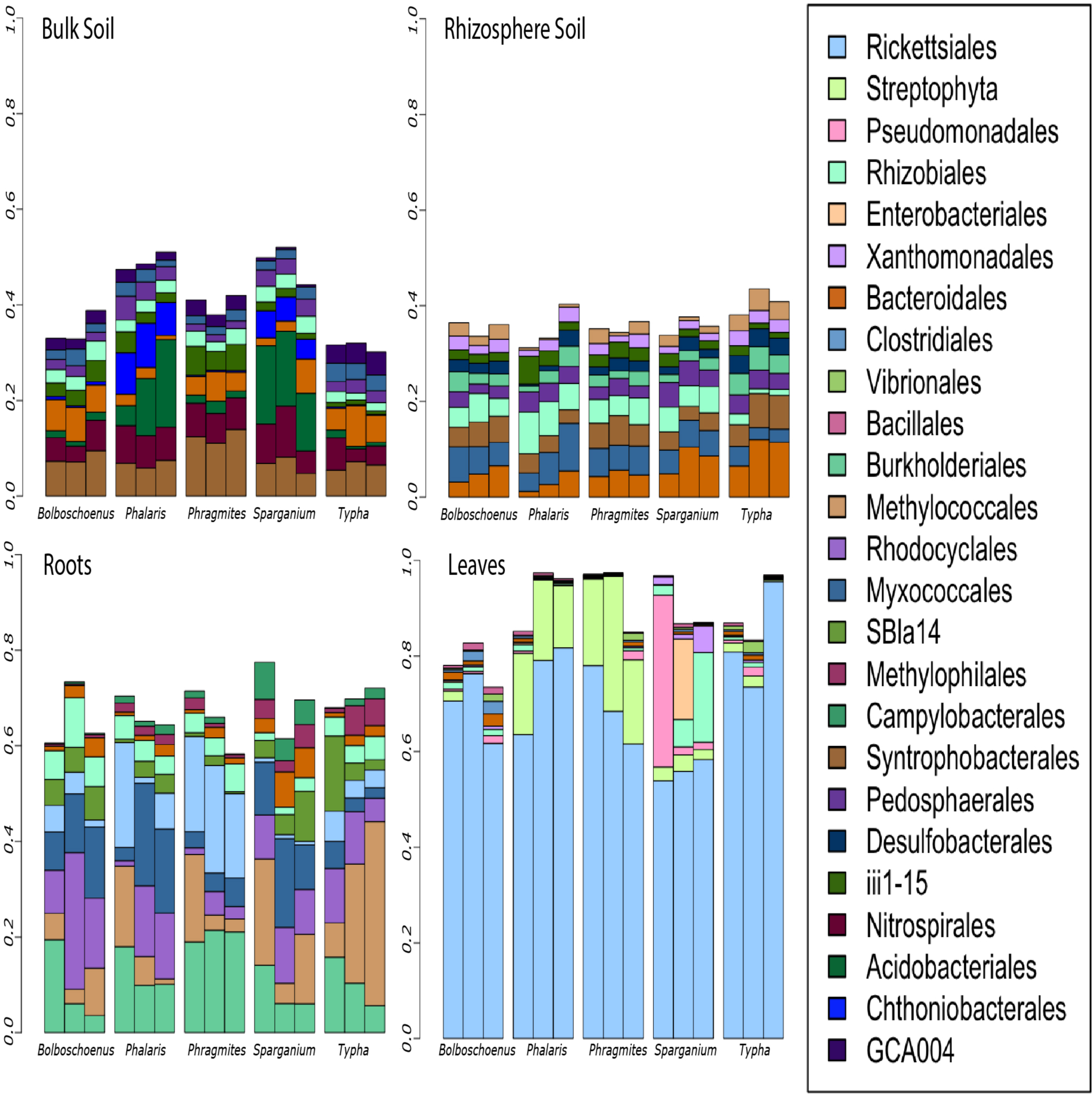
The ten most abundant prokaryote orders for each sample type. Each bar represents the relative abundance of taxa within an individual sample, with three replicate samples per plant genus. Overall alpha-diversity decreases from bulk soil and rhizosphere soil samples through root samples with the least diversity in leaf samples.

Although many microbial orders were present across all sample types, orders which were abundant in soils tended to be more poorly represented in tissues and vice versa (Fig. 2). For example, Syntrophobacterales were abundant in bulk soils (8.0% relative abundance) with decreased relative abundances in rhizosphere soil (5.8%) and low abundance in both roots and leaves (<5.0% each). Similarly, Bacteroidales were the most abundant order in rhizosphere soils (6.1%) with decreased abundance in bulk soils (4.6%) and low abundance in both root and leaf (<2.6%) samples. Burkholderiales were the most abundant order in roots (12.4%) with lower abundances in all other sample types (<1.5%). The Rickettsiales were particularly abundant in leaf and root tissues (70.6% and 7.9%, respectively) but present in low concentrations for all soil samples (<0.19%). Soil and plant tissues, each with unique selective pressures, were host to vastly different microbial communities.

Within the four sample types considered here, both environmental conditions and plant species influenced microbial community composition. Acidobacteriales were more abundant in bulk soils adjacent to the plants in wetland two (*P. arundinacea, S. eurycarpum*, 13.7%% and 14.7%, respectively) than the plants in wetland one (all plant species<2.0%). Meanwhile, the cattails *S. eurycarpum* and *T*. × *glauca* contained higher quantities of Bacteroidales in their rhizosphere soils than other species (8.0% and 10.0% respectively, others <4.9%), despite being in different wetlands. Similarly, the methanotrophic Methylococcales were more abundant in *S. eurycarpum* and *T*. × *glauca* roots than roots of other species (13.7% and 23.5% respectively, others <8.1%). Rickettsiales were more abundant in the true grasses (*P. arundinacea, P. australis*) roots relative to other species (10.2% and 20.0% respectively, others <4.7%). Streptophyta were present in true grass leaves at elevated levels compared to other species (15.7% and 21.3% respectively, others <2.9%). Considerable differences in overall composition at the microbial order level among plant species were also apparent when considering relative abundances of individual OTU.

Across sample types, microbial communities exhibited differences in plant species-specific clustering at the OTU level (Fig. 3). Compared to other sample types, leaf microbial communities were the most consistent across plant species. Similarity in community composition across plant species decreased from leaf through bulk soil communities, where microbial communities were the most distinct by plant species (Fig. 3). While leaf microbial communities were significantly different from other sample types, they were barely distinguishable among plant species (ANOSIM p=0.001, R=0.4296; Permanova p=0.001, r^2^=0.518). Although root microbial communities were still overlapping among plant species, they differed significantly and exhibited distinct clustering (ANOSIM p=0.002, R=0.6459; Permanova p=0.001, r^2^=0.542). Meanwhile, both rhizosphere and bulk soil communities were significantly different and strongly clustered by plant species (ANOSIM p=0.001; R=0.8222 Permanova p=0.001, r^2^=0.624 and ANOSIM p=0.001, R=0.8919; Permanova p=0.001, r^2^=0.001, respectively). Bulk soil samples exhibited the most pronounced clustering by plant species.

**Figure 3.**
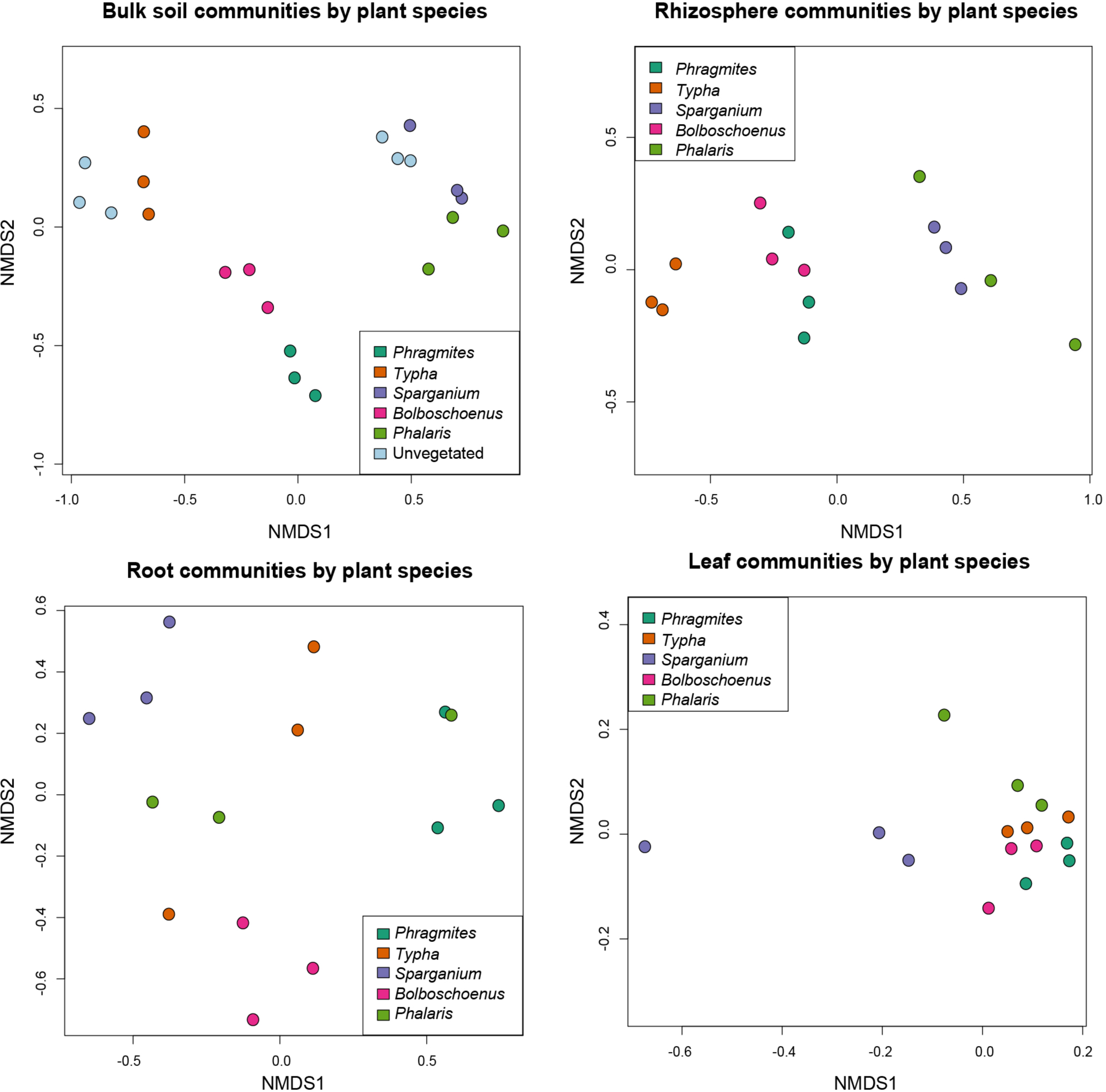
Non-metric dimensional scaling (NMDS) ordination plots representing dissimilarities in microbial community composition at the OTU level within sample types. Clustering according to plant species was least distinctive in plant leaves and most pronounced in soils, particularly bulk soil samples.

For each sample type, dissimilarity in microbial communities was correlated with evolutionary distance among the five host plants. Plant evolutionary history was most closely related to variation in microbial community composition in roots and least in leaves. Twenty percent of the variation in microbial community composition was explained by evolutionary distance among plant species for leaf samples (Mantel p=0.007, r^2^=0.2022). Substantially more variation, 53%, in microbial community composition was explained by evolutionary distance among plant species for root samples (Mantel p=0.001, r^2^=0.5345). Meanwhile, around 40% of the variation in microbial community composition could be explained by evolutionary distance among plant species for both rhizosphere and bulk samples (Mantel p=0.002, r^2^=0.4034; Mantel p=0.001, r^2^=0.4442 respectively).

Overall, differences in microbial community composition were driven both by differences in plant evolutionary history and plant species for each sample types. Each sample type showed different relationships with these factors. Plant species had the weakest effect on leaf communities and the strongest on bulk soil communities. Meanwhile, plant evolutionary history had the strongest effect on root microbial communities and the weakest effect on leaf microbial communities, with an intermediate effect on both soil communities. Microbial community diversity decreased from bulk soil through leaf tissues.

### Methanotroph and Methanogen Populations

*Typha* × *glauca* contained the most methanotrophs across species in every sample type, and its closest evolutionary relative sampled, *S. eurycarpum*, contained the second most methanotrophs in soil samples (Fig.4). *T*. × *glauca* contained significantly larger methanotroph population sizes in bulk soils than *S. eurycarpum*, which contained more than other species (p=0.00456); rhizosphere soils followed the same pattern (p=0.0102). Both leaves and roots of *T*. × *glauca* had larger methanotroph population sizes than other species, although these differences were not significant. *T*. × *glauca* also contained more methanogens in bulk soils than other species (p=0.0237), while *S. eurycarpum* leaves contained significantly more methanogens than either *B. fluviatilis* or *P. arundinacea* (p=0.026). Population sizes of both methanogens and methanotrophs in leaves were at least one order of magnitude smaller than populations of all other samples, and methanogens were at least one order of magnitude smaller in roots than all soil samples. Methanogen population size did not exhibit a clear relationship with plant evolutionary history, but methanotroph population sizes in soils were considerably larger in the cattails *S. eurycarpum* and *T*. × *glauca* than the other plant species.

Methane emissions result from the balance between methane consumption and production. Here we considered the ratio of methanotrophs to methanogens to examine the potential impact of plant species and evolutionary history on methane emissions. The methanotroph to methanogen ratio varied considerably across sample types and plant species (Fig. 4). There was a significant interaction between the effects of plant species and sample type on the ratio of methanotrophs to methanogens (p=0.012). Across plant species, the ratio of methanotrophs to methanogens was similar in bulk soil and unvegetated soil (0.087 and 0.089, respectively); however, this ratio was higher in rhizosphere soil (0.249). Leaves had a greater proportion of methanotrophs to methanogens (0.769) than all soils. However, leaves still contained more methanogens than methanotrophs for each plant species except for *T*. × *glauca*. Roots were the only sample type where methanotroph population sizes were larger than methanogens and this effect was both significant (p<0.0001) and pronounced, with a mean methanotroph to methanogen ratio of 12.2. Populations of both methanotrophs and methanogens were suppressed in bulk and rhizosphere soils compared to unvegetated soil. The methanotroph to methanogen ratio varied considerably across sample types, particularly within roots, and this ratio differed among plant species.

**Figure 4.**
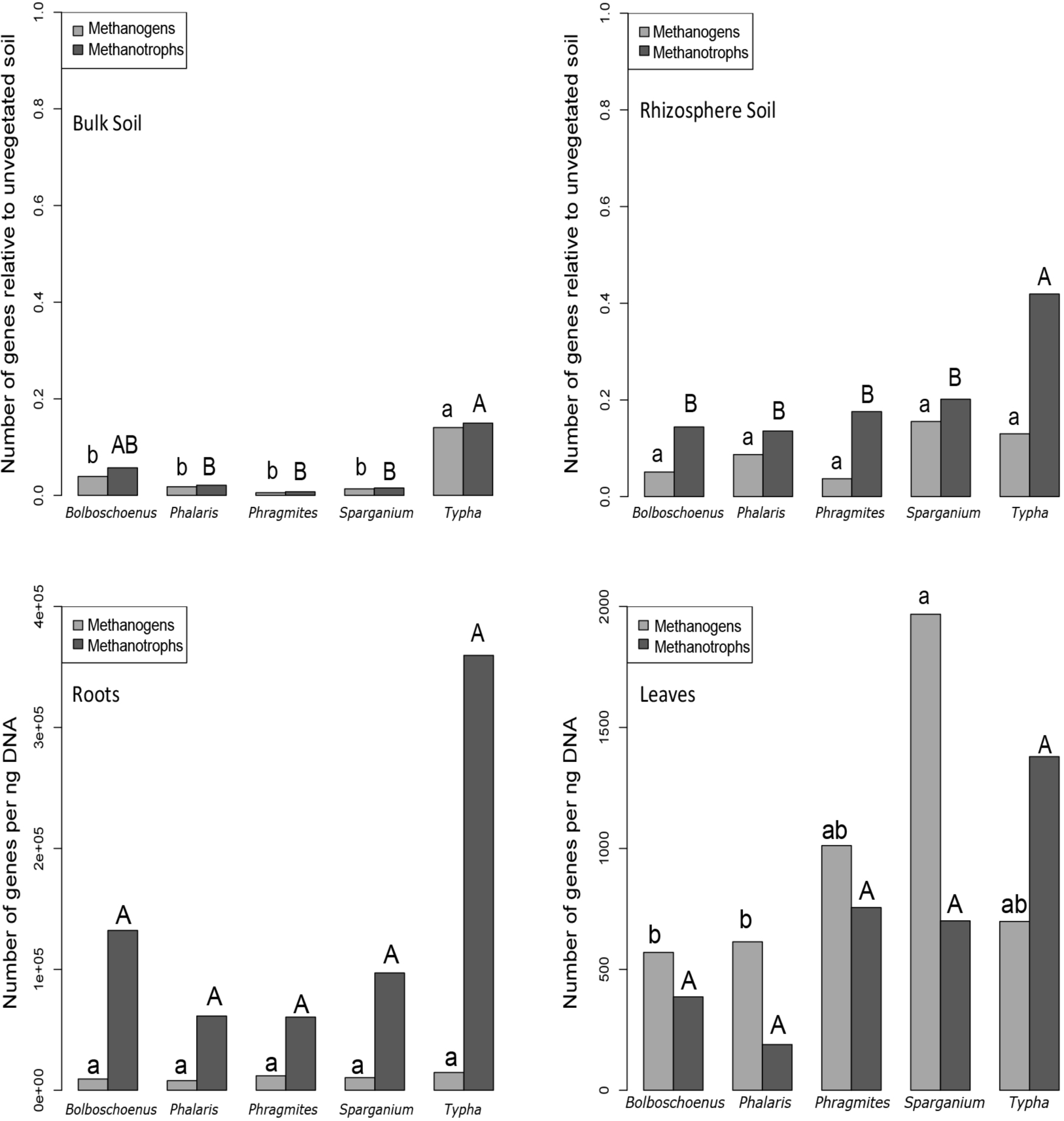
Absolute methanotroph and methanogen abundances normalized to equivalent DNA concentrations for all samples. Soil samples were adjusted in proportion to unvegetated soils by site. Letters above bars denote significant differences between groups according to Tukey’s HSD tests for methanogens and methanotrophs independently. Despite substantial differences in population sizes among roots, no groups were significantly different.

## Discussion

Plant-associated microbial community composition sampled from bulk soil, rhizosphere soil, roots, and leaves differed by both plant species and sample type, with samples distinctly clustered by sample type. While the majority of endophytic taxa were also present in soil samples, soil samples contained a greater number of total and unique OTUs than plant tissues, particularly for the Rickettsiales-dominated leaf communities. This pattern is consistent with the prevailing hypothesis that many root and even leaf endophytes are recruited from soil (Zarraonaindia *et al*. 2015). Additionally, differences in the relative abundance of taxa shared between plant tissues and soil samples were pronounced. Microbial OTUs and orders found to be abundant in soils were present in low abundances within tissues and vice versa, suggesting recruitment and differentiation of communities within plant tissues. For example, Rickettsiales, an order of common plant endophytes (Kunda *et al*. 2018), were abundant in leaf samples for each species and root samples for the true grasses but nearly absent in other sample types. Meanwhile, the methanotrophic order Methylococcales was only abundant in root samples and was most abundant in samples from the Typhaceae. Previous studies have shown that both plant species and environmental conditions influence microbial communities (Berg and Smalla 2009; Gagnon *et al*. 2007), consistent with our finding that wetland plant species substantially influence microbial communities within each sample type.

Plant evolutionary history was related to variation among wetland microbial communities across plant tissues and soil samples. Previous studies have reported dissimilarities in microbial communities relating to phylogenetic distances in the true grass family under greenhouse conditions (Bouffaud *et al*.2014). Our study shows that these differences in plant-associated endophytic and soil microbial communities are related to phylogenetic distance across the grass order in mature, naturally occurring wetland plants. The influence of plant evolutionary history on microbial communities is discernable in the hydrologically similar and physically adjacent wetlands across the plants sampled here. Further work is required to establish the relative importance of evolutionary history when considering additional plant lineages and more distinct wetland sites. Despite the increase in direct control and interaction with the plant, we found that plant evolutionary history explained the least amount of variation in leaf microbial communities compared to other sample types. We attribute this to the lower levels of microbial diversity and richness of unique taxa found in these highly selective environments. While many factors are likely at play, evolutionarily conserved plant morphology and biochemistry, in the forms of aerenchyma (morphology) and root exudates and senesced tissues (biochemistry), are likely mechanisms for the relationship between plant evolutionary history and microbial community composition as redox potential and substrate availability are known drivers of microbial community composition in wetlands (Colmer 2003; Conrad 1996; Megonigal 2004).

The population sizes of methanotrophs and methanogens within these plant-associated microbial communities differ by sample type and plant species, offering one putative explanation for why some variation in methane emissions can be explained by plant species composition (Wang and Han 2005; Yavitt and Knapp 1998). Factors known to influence methane-cycling microbes, such as redox potential which can be influenced by both aerenchyma flow rates and organic matter deposition, vary considerably across plant species and are phylogenetically conserved (Jung *et al*. 2008; Visser *et al*. 2000; Reich *et al*. 2003). Despite this, methanogen population sizes did not exhibit a clear relationship with plant evolutionary history and were relatively stable across samples in agreement with previous work (Goldberger 2012). Methanotrophs, on the other hand, showed increases in population size within *T*. × *glauca*.

The combination of highly aerobic roots and high gas diffusivity through *Typha* (Brix *et al*. 1992; Konnerup *et al*. 2011), particularly for the deeply submerged *T*. × *glauca*, may explain the exceptionally large methanotroph population sizes found in particular plants. A similarly high ratio was not observed in its sister taxon, *S. eurycarpum*, despite a high abundance of Methylococcales in roots, more than the other species sampled. The particularly large methanotroph to methanogen ratio within *T*. × *glauca* roots, in combination with closed stomates at night, may explain previous observations of high carbon dioxide concentrations within *Typha* leaves at night (Constable *et al*. 1992). *Typha latifolia* has been shown to indirectly consume larger quantities of methane through root and rhizome associated methanotrophs than wetland sedges and true grasses (King 1994), indicating that *Typha* species other than *T*. × *glauca* may increase methane oxidation in wetlands. This methane utilization may provide a fitness advantage for *Typha* species as many wetland plants have been shown to utilize the carbon dioxide produced by methanotrophs for photosynthesis (Raghoebarsing *et al*. 2005). The impacts these microbes have on their plant hosts’ fitness, particularly regarding the large population sizes of methanotrophs associated with *T*. × *glauca* roots, remains unknown but may partially explain why *T*. × *glauca* can often successfully invade deeply flooded marsh habitats.

Our finding of an increased similarity of microbial community composition with more closely related species suggests that plant evolutionary history may provide additional insight into observed variability in wetland methane emissions. For example, the sampled plant taxa in the Typhaceae, *S. eurycarpum* and *T*. × *glauca*, harbor high population densities of methanotroph orders, so wetlands dominated by species within the Typhaceae may have lower methane emissions than true grass and sedge-dominated wetlands. Notably, the exceptionally high methanotroph population sizes in *T*. × *glauca* roots measured here, in agreement with previous studies on methanotroph populations in *T. latifolia* roots and rhizomes (King 1994), suggests that wetlands inhabited by *Typha* species may have particularly low methane emissions. This conclusion runs counter to predictions based solely on environmental conditions which would suggest that the more deeply submerged wetlands in which *Typha* dominate would have the greatest potential for methanogenesis and least for methane oxidation. In addition to providing a potential mechanism underlying the relationship between plant community composition and the ecosystem processes of methanogenesis and methanotrophy in wetlands, the relationships between wetland plant species and their endophytic and soil microbiomes found here potentially affect plant fitness and species distribution patterns through changes in carbon dioxide availability within plant tissues.

## Acknowledgements

We would like to thank Dion Antonopoulos for his assistance with obtaining the gBlocks used in qPCR for both methanotroph and methanogen amplification. Theodore Flynn provided initial guidance in processing the Mi-Seq output data. The Summer Undergraduate Laboratory Intern (SULI) program, hosted by Argonne National Laboratory, funded Marisa (Austin) Szubryt during the project’s conception and three other student interns who assisted with project development and review during its early stages: Maria Pia Ramos, Lauren Sinclair Johnson, and Korbibian Thalhammer. We wish to thank the Society of Wetland Scientists for providing an undergraduate Student Research Grant worth $948 to Marisa Szubryt in spring of 2017. This research is part of the Wetland Hydrobiogeochemistry Scientific Focus Area (SFA) at Argonne National Laboratory supported by the Subsurface Biogeochemical Research Program, Office of Biological and Environmental Research, Office of Science, U.S. Department of Energy (DOE), under contract DE-AC02-06CH11357.

