## Supplemental Figures for "Wetland plant evolutionary history influences soil and endophyte microbial community composition"

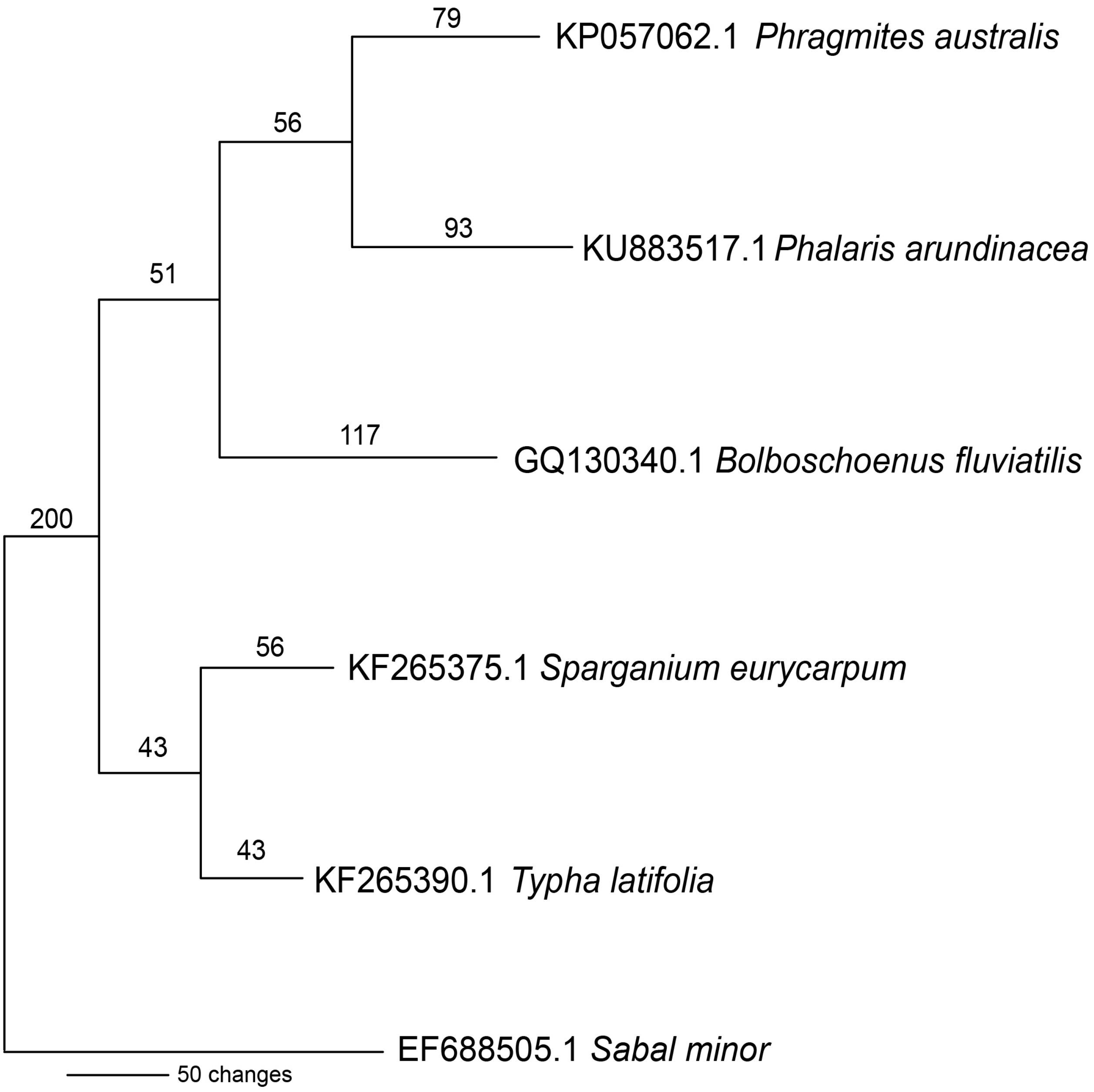


**Supplemental Figure 1**. Internal transcribed spacer phylogeny of host plant taxa with the palm *Sabal minor* of Arecales as an outgroup to these Poales taxa. Numbers above each branch indicate branch length based on PAUP parsimony heuristic searches.

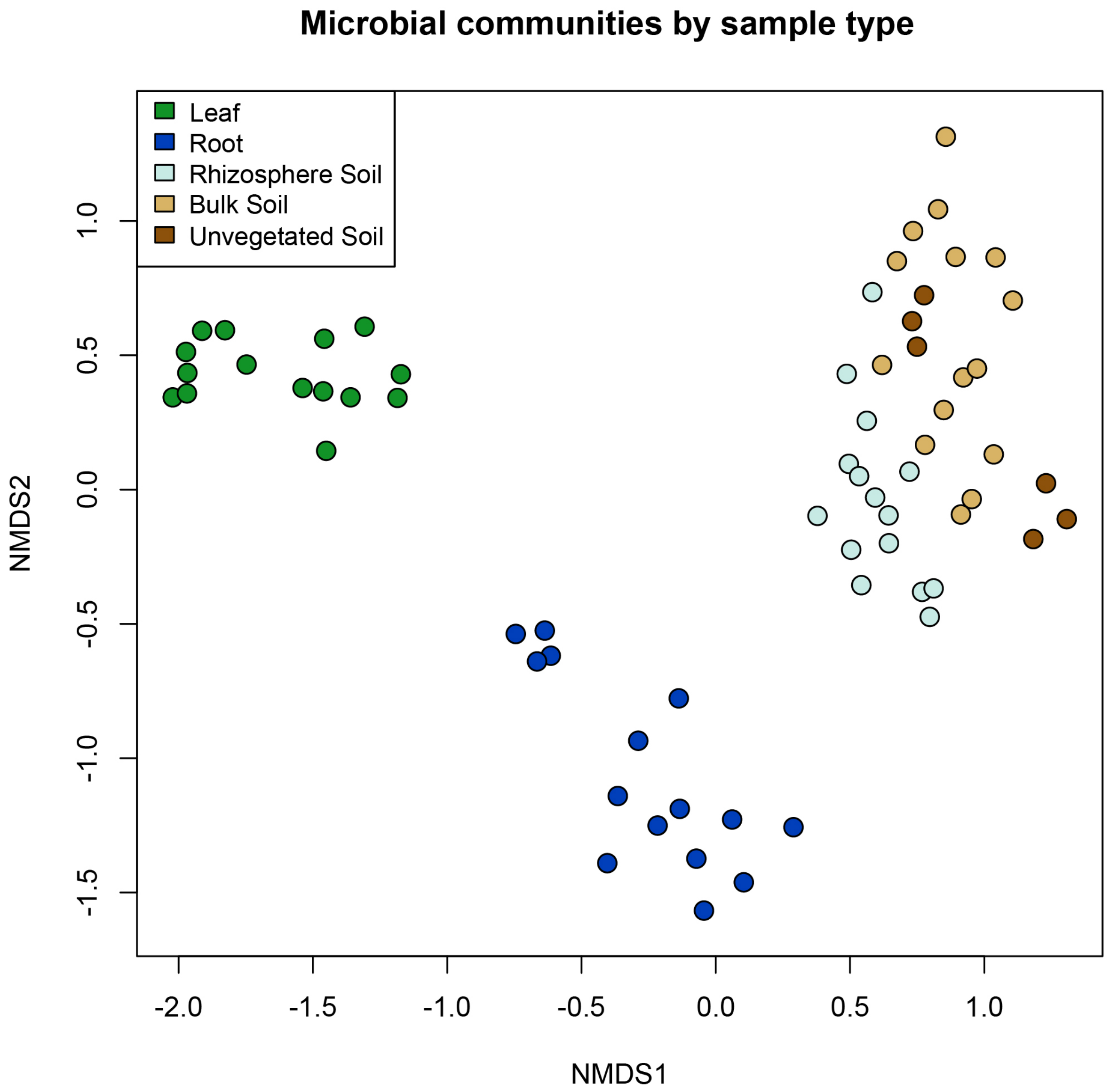


**Supplemental Figure 2.** Non-metric multidimensional scaling (NMDS) ordinations of every sample showing distinct clustering by sample type.
